# Analysis of interphase node proteins in fission yeast by quantitative and super resolution fluorescence microscopy

**DOI:** 10.1101/137794

**Authors:** Matthew Akamatsu, Yu Lin, Joerg Bewersdorf, Thomas D. Pollard

## Abstract

We used quantitative confocal microscopy and FPALM super resolution microscopy of live fission yeast to investigate the structures and assembly of two types of interphase nodes, multiprotein complexes associated with the plasma membrane that merge together and mature into the precursors of the cytokinetic contractile ring. During the long G2 phase of the cell cycle seven different interphase node proteins maintain constant concentrations as they accumulate in proportion to cell volume. During mitosis the total numbers of type 1 node proteins (cell cycle kinases Cdr1p, Cdr2p, Wee1p, and anillin Mid1p) are constant even when the nodes disassemble. Quantitative measurements provide strong evidence that both types of nodes have defined sizes and numbers of constituent proteins, as observed for cytokinesis nodes. Type 1 nodes assemble in two phases, a burst at the end of mitosis, followed by steady increase during interphase to double the initial number. Type 2 nodes containing Blt1p, Rho-GEF Gef2p, and kinesin Klp8p remain intact throughout the cell cycle and are constituents of the contractile ring. They are released from the contractile ring as it disassembles and then associate with type 1 nodes around the equator of the cell during interphase.

**Highlight summary:** FPALM super resolution microscopy and quantitative confocal microscopy reveal that interphase nodes, the precursors to the fission yeast cytokinetic contractile ring, are discrete unitary structures with defined sizes and ratios of component proteins. Type 1 nodes disassemble during mitosis, but type 2 nodes remain intact throughout the cell cycle.

## INTRODUCTION

Fission yeast divide by cytokinesis, when constriction of a contractile ring made of actin and myosin (Stachowiak *et al*., 2014) and growth of the extracellular septum (Proctor *et al*., 2012) form a furrow at the equator of the cell. The contractile ring forms from clusters of cytokinesis proteins called nodes that are associated with the plasma membrane around the middle of the cell (Vavylonis *et al*., 2008). Cytokinesis nodes have stoichiometric proportions of constituent proteins (Wu and Pollard, 2005; Laporte *et al*., 2011). Super resolution measurements (Laplante *et al*., 2016b) and two-color confocal distance measurements (Laporte *et al*., 2011) show that cytokinesis nodes have a defined organization, with myosin-II tails anchored to a compact base near the plasma membrane containing anillin Mid1p and myosin-II heads projecting into the cytoplasm. Mid1p serves as a scaffold for other cytokinesis proteins (Paoletti and Chang, 2000; Celton-Morizur *et al*., 2004; Wu *et al*., 2006; Almonacid *et al*., 2009; Laporte *et al*., 2011; Saha and Pollard, 2012b).

During interphase two types of nodes composed of different proteins appear in distinct regions of the cell and merge to form cytokinesis nodes (Akamatsu *et al*., 2014). Stationary type 1 nodes containing cell cycle kinases Cdr1p, Cdr2p and Wee1p form around the middle of cells early during interphase and accumulate anillin Mid1p from the nucleus (Martin and Berthelot-Grosjean, 2009; Moseley *et al*., 2009). The type 2 node proteins Blt1p, Gef2p and Nod1p are concentrated in contractile rings and emerge as discrete, punctate structures as the ring disperses (Moseley *et al*., 2009; Zhu *et al*., 2013; Akamatsu *et al*., 2014). During interphase type 2 nodes diffuse along the cell cortex until they are captured by stationary type 1 nodes around the equator (Akamatsu *et al*., 2014). The combined nodes subsequently accumulate cytokinesis proteins early in mitosis (Moseley *et al*., 2009; Saha and Pollard, 2012) to form cytokinesis nodes. Animal cells may also use node-like clusters of contractile ring proteins as precursors for contractile rings (Straight *et al*., 2003; Werner *et al*., 2007; Hickson and O’Farrell, 2008; Zhou and Wang, 2008; Wollrab *et al*., 2016; Henson *et al*., 2017).

Previous papers left open four important questions about interphase nodes. First, the wide range of fluorescence intensities of interphase nodes marked with Cdr2p, Mid1p, Gef2p, or Blt1p (Laporte *et al*., 2011; Ye *et al*., 2012; Zhu *et al*., 2013; Pan *et al*., 2014) raised the question of whether interphase nodes are defined units or if they might be amorphous protein aggregates. It was uncertain whether the compositions of interphase nodes are heterogeneous, or if the presence of variable numbers of unitary structures was obscured by the limited resolution of the confocal microscope, or both.

Second, the numbers of nodes during interphase was uncertain. Three studies using confocal microscopy agreed that type 1 nodes marked by Cdr2p-GFP double in number during interphase (Deng and Moseley, 2013; Bhatia et al., 2014; Pan et al., 2014). However, the reported numbers of type 1 nodes varied from 10 to 20 in a mid-focal plane (Bhatia *et al*., 2014) to 25 to 40 (Pan *et al*., 2014) or ∼52 to 106 in whole cells (Deng and Moseley, 2013).

Third, incomplete information about the cellular concentrations of interphase node proteins (especially those in type 2 nodes) left us wondering if any of their concentrations change during the cell cycle. Quantitative confocal microscopy was used to estimate the numbers of interphase node proteins including Mid1p (Wu and Pollard, 2005; Coffman *et al*., 2011; Laporte *et al*., 2011), Gef2p, Nod1p (Zhu *et al*., 2013) and Cdr2p (Pan *et al*., 2014) in whole cells and interphase nodes. Mass spectrometry showed that the numbers of Cdr2p, Blt1p and Klp8p were constant around the time of mitosis, but no measurements were made for the other ∼80% of the cell cycle during interphase (Carpy *et al*., 2014). No quantitative measurements were available for Klp8p, Cdr1p or Wee1p.

Fourth, Blt1p, the presumed scaffold for type 2 nodes, is incorporated into contractile rings, but nodes are not resolved within rings by confocal microscopy (Moseley *et al*., 2009; Goss *et al*., 2014). Therefore, we did not know if nodes remain intact or if Blt1p disperses in contractile rings.

We used quantitative confocal microscopy and high-speed FPALM (fluorescence photoactivation localization microscopy) (Huang *et al*., 2013) of live fission yeast cells to address each of these questions. We investigated the organization and stoichiometry of proteins in interphase nodes across the cell cycle. Both types of measurements suggest that interphase nodes are unitary structures that increase in number over the cell cycle. The observations also confirmed our speculation (Akamatsu *et al*., 2014) that type 2 nodes marked by Blt1p persist as discrete structures within the contractile ring.

## RESULTS

### Interphase node proteins accumulate in proportion to cell volume

We measured the total numbers per cell of seven interphase node proteins tagged with GFP or mEGFP at the N- or C-terminus by quantitative confocal microscopy (Figure 1, Table 1). Six of these proteins were expressed from the genome under the control of their native promoters. mEGFP-Gef2p was overexpressed due to a second promoter upstream of the resistance gene (Zhu *et al*., 2013). These tagged strains grew normally (Figure S1). We examined populations of asynchronous cells and used cell lengths to estimate cell cycle time.

**Figure 1.**
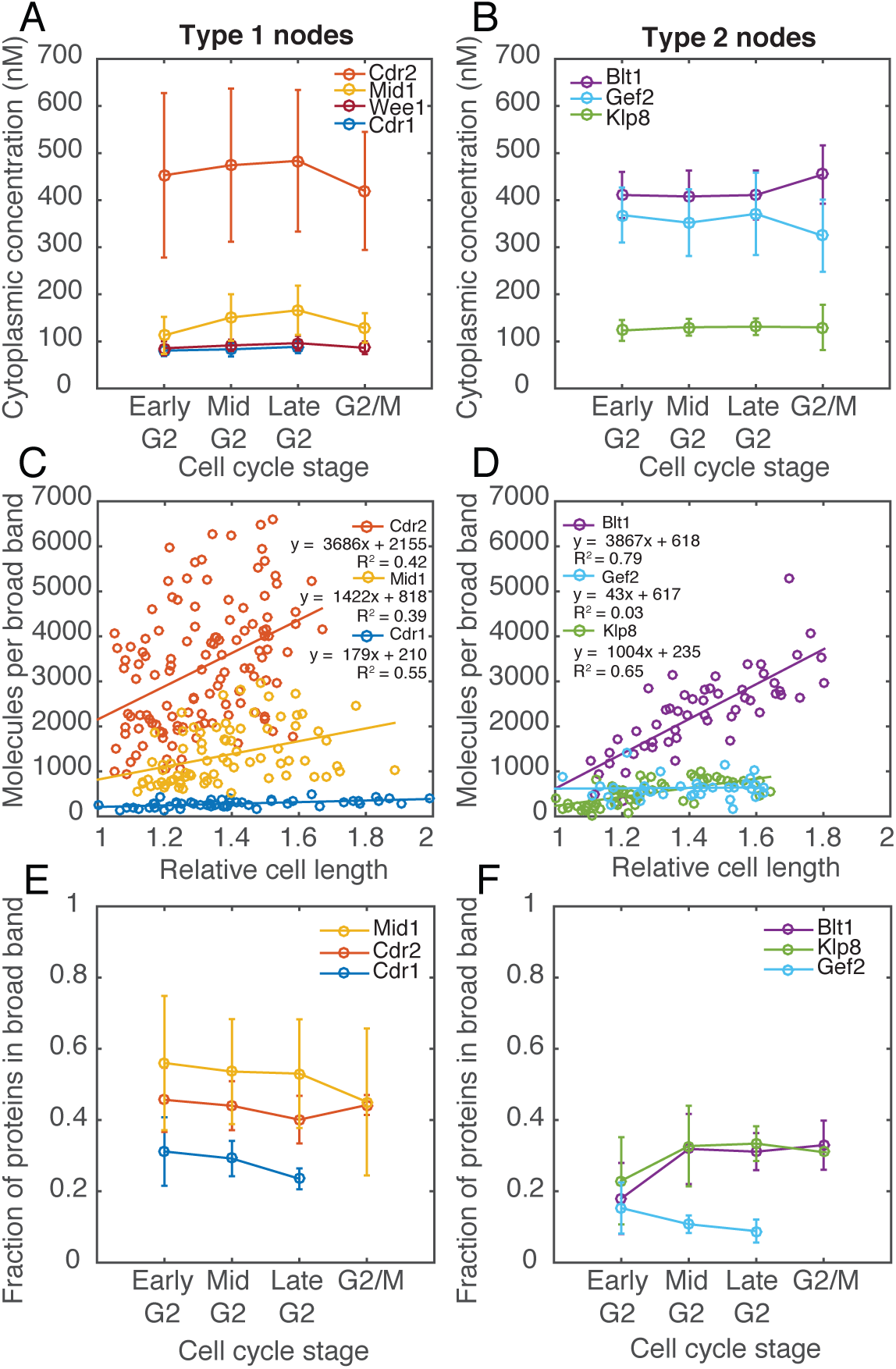
Global and local numbers of molecules of interphase node proteins over the cell cycle measured by quantitative fluorescence microscopy as explained in the methods section. Values in A, B, E, F are means ± SD. (A, B) Plots of the concentrations of (A) type 1 and (B) type 2 interphase node proteins over the cell cycle. Cells were classified into interphase stages by cell length. (C, D) Plots of numbers of molecules in the broad band of nodes over the cell cycle for (C) type 1 and (D) type 2 interphase node proteins. The X-axis, relative cell length, is defined as the cell length scaled relative to the smallest (1) and largest (2) cell in the population. Extrapolation of linear fits to X = 1 give the average number of molecules at the cell equator at cell birth. Cells expressing Cdr1p-3GFP were shorter than wild type cells (Martin and Berthelot Grosjean, 2009; Moseley *et al*., 2009) so their lengths were scaled independently. (E, F) Plots of the fractions of each interphase node protein in the broad band of nodes over the cell cycle for (E) type 1 and (F) type 2 nodes. Cells were classified into interphase stages by cell length.

**Table 1.**
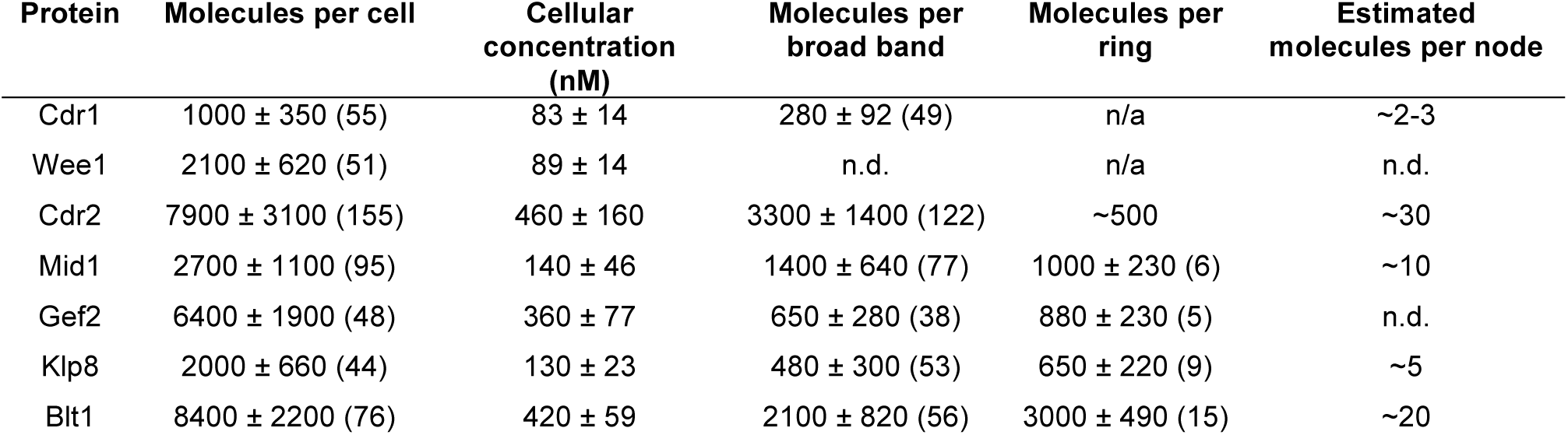
Measurements by quantitative confocal fluorescence microscopy of the numbers of molecules of interphase node proteins per cell, in the broad band and per node (number of molecules per broad band of nodes divided by the number of nodes). Measurements of molecules in the contractile ring were made for fully formed (continuous) unconstricting rings. Values ± SD with number of cells measured in parentheses. n.d., not determined; n/a, not applicable.

The cellular concentrations of the four type 1 node proteins (Cdr1p, Cdr2p, Wee1p and Mid1p) and three type 2 node proteins (Gef2p, Blt1p and Klp8p) were constant during the cell cycle (Figure 1, A and B), that is their total numbers per cell increased in proportion to cell size. The scaffold proteins Cdr2p and Blt1p are present at higher concentrations than the other node proteins.

Our data confirm previous fluorescence intensity measurements on Mid1p (Wu and Pollard, 2005) using a range of calibration standards and on Cdr2p (Pan *et al*., 2014) using the ratio of Cdr2p-GFP to Rlc1p-GFP and the concentration of Rlc1p-GFP (Wu and Pollard, 2005). Our numbers also agree within a factor of two with measurements by mass spectrometry of Cdr2p, Blt1p and Klp8p during the G2-S interval of the cell cycle (Carpy *et al*., 2014). That study did not report the numbers of these node proteins for the vast majority of interphase, about 80% of the cell cycle and the time of interest for our study. Carpy et al. examined cells arrested at G2/M for 5 h, which produced a heterogeneous population of abnormally large cells. They divided all of their numbers by 3.5, an estimate of the average difference in size of their cell population compared and wild type cells. They and Zhu et al. (2013) reported ∼2000 Gef2p molecules at the end of G2, so Gef2p was over expressed by about 3.5-fold in our cells. Other counts of cytokinesis proteins by mass spectrometry (Marguerat *et al*., 2012) differ from fluorescence measurements, even by more than an order of magnitude (Coffman and Wu, 2014).

### Interphase node proteins accumulate around the equator during interphase

Owing to their accumulation in nodes, the total numbers of type 1 node proteins increased linearly in the broad band around the equator during interphase (Figure 1, C and D). These measurements are consistent with previous qualitative observations (Deng and Moseley, 2013; Pan *et al*., 2014). Type 1 nodes form around the equator before cell separation (Paoletti and Chang, 2000; Morrell *et al*., 2004; Martin and Berthelot-Grosjean, 2009; Moseley *et al*., 2009; Akamatsu *et al*., 2014), so the numbers of type 1 node proteins in the broad band are already high in short early G2 cells. Linear regression fits to the counts of molecules vs. cell length allowed us to determine the mean number of molecules in the broad band of nodes (y-intercepts) in early interphase: ∼2200 Cdr2p, ∼800 Mid1p, and 200 Cdr1p. The numbers of all three proteins approximately doubled during interphase (Figure 1C) before Cdr2p and Cdr1p dispersed from nodes during mitosis. When expressed at endogenous concentrations the kinase GFP-Wee1p was present in interphase nodes (Figure S2B), but the low signal and its presence at the spindle pole body (SPB) precluded accurate measurements of its numbers in the equatorial broad band. The fraction of each type 1 node protein in the broad band of nodes remained relatively constant throughout interphase (30-50% of the total cellular pool) (Figure 1E).

Type 2 nodes began the cell cycle concentrated at the new poles created by cytokinesis (Moseley *et al*., 2009; Saha and Pollard, 2012; Akamatsu *et al*., 2014; Goss *et al*., 2014), and their numbers around the equator were low. During interphase, type 2 nodes moved to the cell equator (Akamatsu *et al*., 2014) where the total numbers of Blt1p and Klp8p increased 4-5 fold (Figure 1D). From early G2 to mid G2 the fraction of Blt1p and Klp8p located in the broad band of nodes increased from 20% to 30% (p<0.01) (Figure 1F). Once type 1 and 2 nodes merged around the equator (Akamatsu *et al*., 2014), the ratios of their constituent proteins in nodes remained fairly constant (Table 2). Overexpressed mEGFP-Gef2p did not increase around the equator during interphase, most likely because most of the signal was in the cytoplasm and was included in the measurements of the broad band.

**Table 2.** Ratios of the numbers of molecules per node at different stages of interphase relative to the least abundant protein measured (Cdr1p). Values ± SD with number of cells measured in parentheses. n.d., not determined

| Protein | Early G2 | Mid G2 | Late G2 | G2/M | Late interphase |
| --- | --- | --- | --- | --- | --- |
| **Cdr1** | 1 (21) | 1 (19) | 1 (9) | n.d. | 1 |
| **Cdr2** | 14 ± 6 (44) | 12.5 ± 5 (44) | 12 ± 5 (32) | 12 ± 2 (2) | 12 |
| **Mid1** | 4.5 ± 6 (17) | 5 ± 2 (36) | 5.5 ± 2 (17) | 3.5 ± 2 (8) | 5 |
| **Gef2** | 4 ± 2.5 (7) | 2 ± 0.5 (14) | 2 ± 0.5 (17) | n.d. | n.d. |
| **Klp8** | 1.5 ± 0.5 (27) | 2 ± 0.5 (14) | 2 ± 0.5 (12) | 1 ± 0 (1) | 2 |
| **Blt1** | 5 ± 2.5 (10) | 8 ± 2 (17) | 8 ± 2 (17) | 8 ± 2 (10) | 8 |

We calculated the ratios of molecules in the broad band during late interphase relative to the least abundant protein measured: 1 Cdr1p: ∼12 Cdr2p, 5 Mid1p, 2 Klp8p, and 8 Blt1p (Table 1). These ratios changed little between mid G2 and G2/M (Table 2).

### Numbers of proteins in interphase nodes are quantized

We used quantitative confocal microscopy to measure the fluorescence intensities of individual nodes of interphase fission yeast cells expressing one of seven interphase node proteins tagged with mEGFP at the N- or C-terminus. We collected z-series of confocal images and measured the fluorescence intensities in 5 confocal slices separated by 260 nm (Figure 2A). The fluorescence intensities of individual interphase nodes were stable during 2.5 min of observation at intervals of 1 or 2 s after correcting for the overall photobleaching of the cells (Figure S3). The two presumed scaffold proteins, Cdr2p and Blt1p, are the most abundant proteins in the two types of nodes.

**Figure 2:**
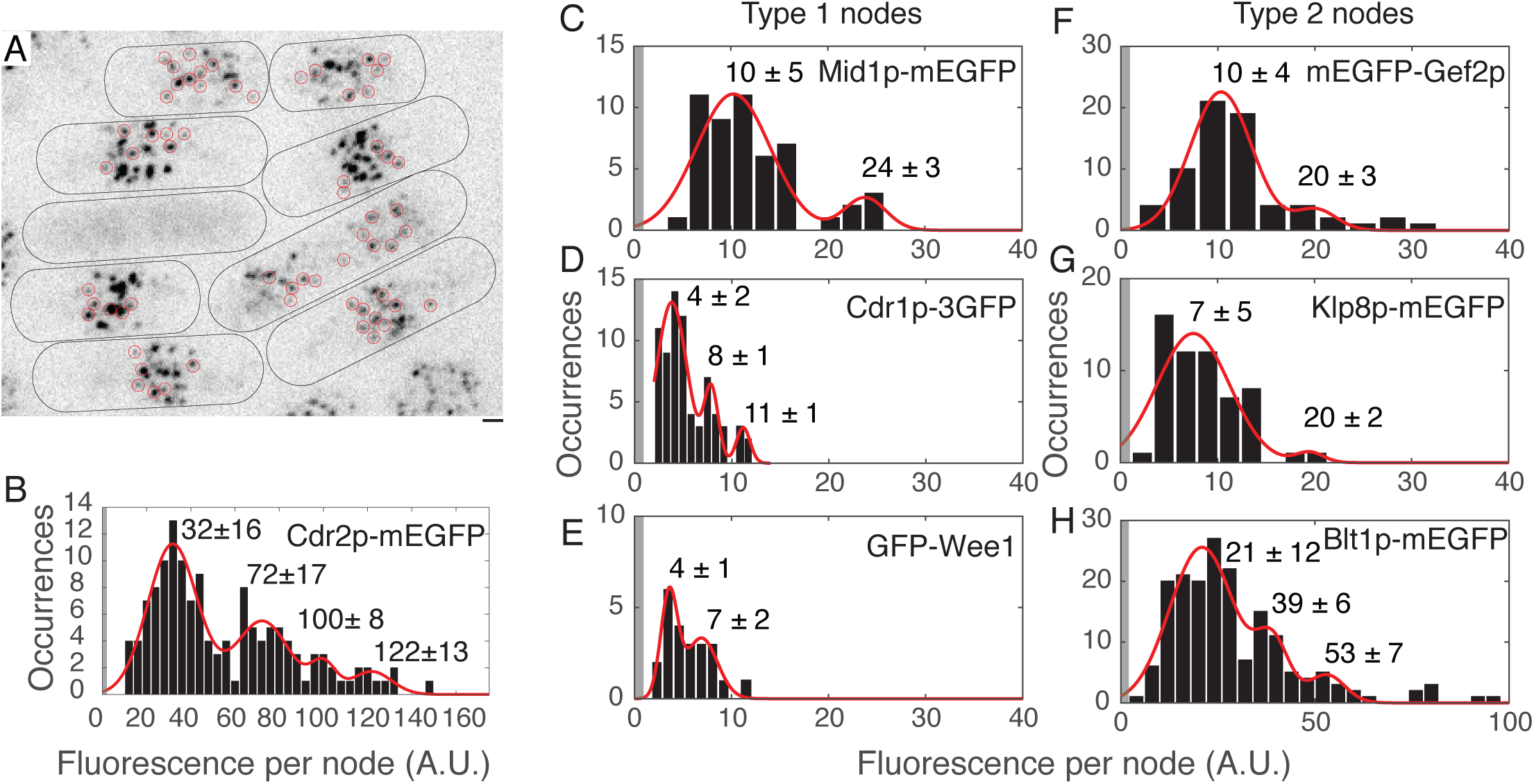
Fluorescence intensities of individual type 1 and 2 interphase nodes. (A) Field of cells expressing Cdr2p-mEGFP. Reverse contrast fluorescence micrograph of a sum projection of 5 slices closest to coverslip. Red circles are node regions selected for fluorescence measurements (see methods). Black lines outline cells. Scale bar 1 μm. (B-H) Histograms of fluorescence intensity per node. The continuous curves are fits of multiple Gaussian distributions to the data. Peak values are means ± SD from the fits. Shaded region is background fluorescence intensity + 1 SD. n = (B) 138 spots in 33 cells; (C) 23 spots in 4 cells; (D) 72 spots in 20 cells; (E) 23 spots in 11 cells; (F) 68 spots in 18 cells; (G) 58 spots in 17 cells; 177 spots in 29 cells.

Both type 1 and 2 interphase nodes varied widely in fluorescence intensities and size in the confocal microscope (Figure 2A, Coffman *et al*., 2011; Laporte *et al*., 2011; Ye *et al*., 2012; Zhu *et al*., 2013; Pan *et al*., 2014). The large, bright, heterogeneous nodes attracted the eye, but most nodes had low intensities and uniform sizes. We used two strategies to measure fluorescence intensity in these nodes. The analysis in Figure 2 was restricted to small nodes that were well separated from other nodes and with their fluorescence included entirely in 5 z-sections. We measured the fluorescence in regions 7 pixels (∼583 nm) in diameter encircling these small nodes.

The fluorescence intensities of the interphase node proteins were binned into peaks corresponding to multiples of a unitary value (Figure 2, B-H). Nodes with the unitary fluorescence were the most numerous. For example, in the sampled population of nodes marked with Cdr2p 57% were unitary, 28% were binary and the remainder had higher fluorescence intensities. Thus some nodes are located so close together that we count them as one node with a multiple of the unitary fluorescence. These fluorescence intensity histograms were better fit by multiple Gaussian distributions (Figure 2) than by continuous log-normal distributions (Figure S2, H-N; Table S3). In cells co-expressing Cdr2p-mCherry and Cdr1p-3GFP, the fluorescence intensities per node were correlated in the two channels (Figure S2, C-E).

The above analysis excluded the brightest nodes, which represented at least half of the total nodes. Using larger regions of interest to examine a sample of bright nodes that were well separated from each other, we found that their fluorescence intensities were up to 10 times the unitary fluorescence (Figure S2, A-C). The fluorescence per unitary node did not change as a function of cell length (Figure S4, Pan *et al*., 2014).

We conclude that nodes are discrete structures with stoichiometric ratios of proteins, but many nodes are too close together to be resolved by confocal microscopy even during interphase. Therefore, we turned to super resolution microscopy to test this hypothesis.

### Super resolution microscopy of interphase nodes

We used the photoconvertable fluorescent protein mEOS3.2 (Zhang *et al*., 2012) to tag the C-termini of Cdr2p and Blt1p in the genome, so the native promoter controlled the expression of the fusion proteins. Both strains grew normally (Figure S1). Figure S5 explains how we acquired FPALM images of these cells at 200 frames per second using laser intensities that photoconverted, imaged and bleached most of the tagged proteins in a field within 50 s. The localization precision was ∼35 nm (Figure S5C). We imaged nodes from two perspectives, either by focusing in the middle of the cell to obtain side views of nodes along the sides of cells (Figures 3, A-C and 4, A B E and F) or by focusing on the cell surface to obtain face views (Figures 3, D-G and 4, D and F-I).

High speed FPALM of cells expressing tagged interphase node proteins showed that both Cdr2p-mEOS3.2 and Blt1p-mEOS3.2 concentrated in discrete structures less than 100 nm in diameter (Figures 3, A-C and 4, A B E and F). The following sections trace the life histories, numbers, locations and surface densities of interphase nodes throughout the cell cycle.

### Super resolution microscopy of type 1 interphase nodes

After being dispersed in the cytoplasm during mitosis (Figures 2A and 3Bj), Cdr2p-mEOS3.2 reappeared in discrete, punctate structures less than 100 nm in diameter on the inside of the plasma membrane around the nuclei of both daughter cells (Figure 3A, a and b). Throughout interphase Cdr2p-mEOS3.2 remained localized to these discrete structures distributed in a broad band around the cell equator. These images of type 1 nodes were sharp, because type 1 nodes have low diffusion coefficients (Akamatsu *et al*., 2014) and moved less than the precision of the imaging method during the time of acquisition. These punctate structures appeared similar in side views (Figure 3, A-C) and face views (Figure 3E). The appearance of Cdr2p-mEOS3.2 in these new nodes was indistinguishable from type 1 nodes later during interphase, so we did not detect any intermediate forms.

**Figure 3:**
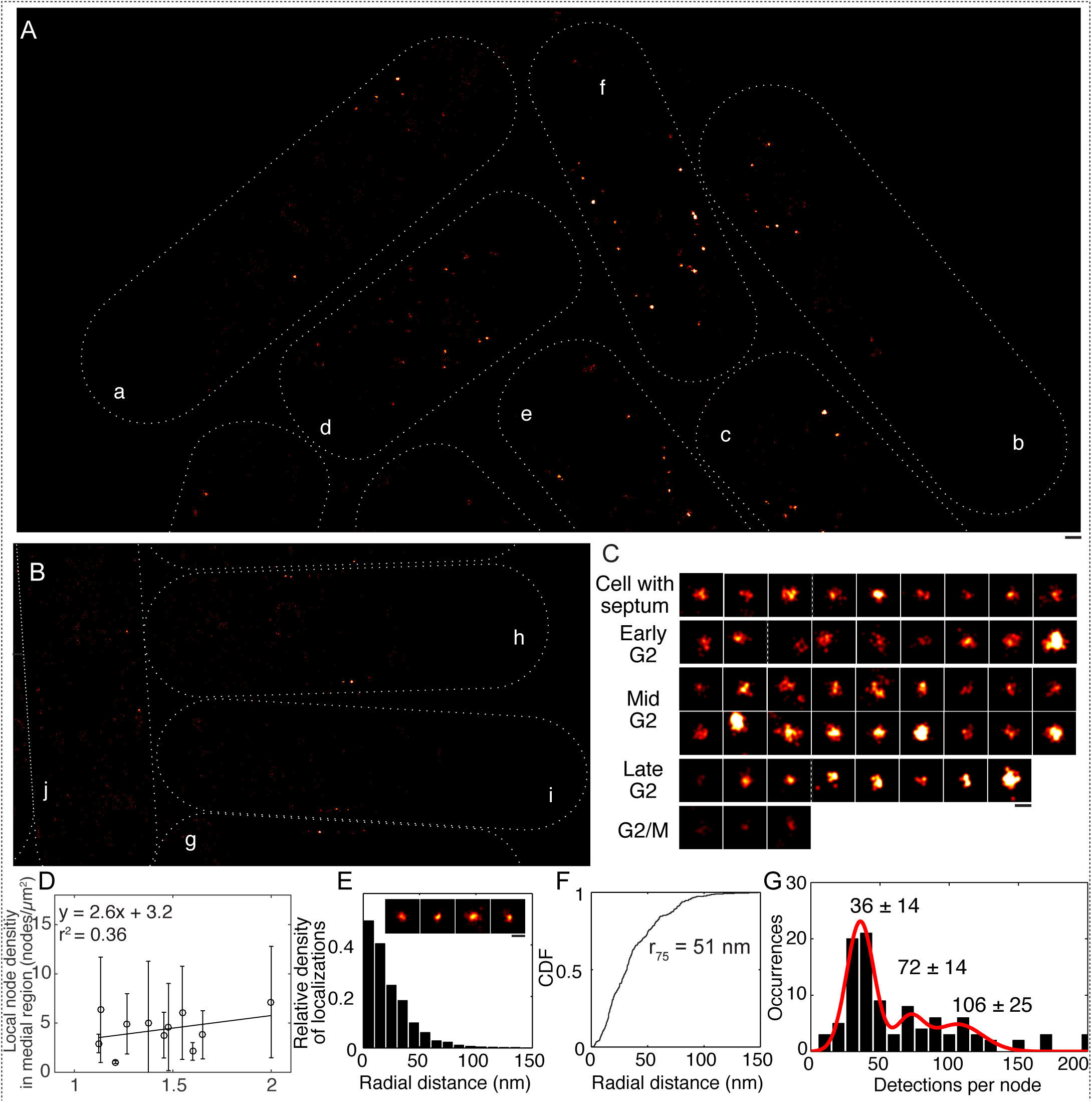
High speed FPALM of cells expressing Cdr2p-mEOS3.2. (A, B) Images displayed as Gaussian kernel density heat maps in focal planes part way between a medial longitudinal section and the cell surface, which captures nodes near the surface. Cells are labeled a through i in the order of cell cycle stage based on length and the presence of a septum: a, b = cells with septa; c, d = early G2; e, f = mid G2; g = late G2; h = G2/M; i = mitosis. White lines mark cell perimeters. Bar is 400 nm. (C) Images of individual nodes of cells marked in A and B with lower contrast. See methods for details on cell classification. Dashed white lines separate nodes from different cells. Bar is 100 nm. (D) Surface densities of interphase nodes in a zone 1.6 μm wide centered on the equator across the cell cycle. Densities were determined by Voronoi tessellation (see Figure S6). The sample was 122 nodes in 11 cells in three fields. Line is a linear fit. (E and F) Analysis of the spatial distribution of Cdr2p-mEOS3.2 in face views of nodes with fewer than 55 detections (approximately the n = 1 peak in G). (E) Histograms of the radial density distribution of mEOS3.2 detections from the center of each node. Inset: Gaussian kernel density heat maps of detections in individual nodes (face views). Bar is 100 nm. (F) Cumulative distribution plots of radial density of detections in nodes marked by Cdr2p-mEOS3.2. The 75th percentile of detection radial distances is reported in the figure. CDF, cumulative distribution function. (G) Histogram of the numbers of FPALM detections per node for face view of Cdr2p-mEOS3.2 nodes. The continuous curves are fits of multiple Gaussian distributions to the data with the peak numbers of detections indicated. Values reported are means ± SD from the fits. N = 92 spots.

The numbers of localized detections of Cdr2p per node covered a wide range, but clustered in discrete peaks corresponding to multiples of a unitary number of 36 detections per node (Figure 3G), similar to our confocal microscopy observations (Figure S4I and Table S3). In the sampled population of nodes marked with Cdr2p (all discrete spots with Gaussian distributions of detections) 58% were unitary, 17% were binary and the remainder had higher numbers of detections. Thus many nodes were located so close to neighbors that they were not resolved even by super resolution microscopy. The numbers of localized detections per node did not change as a function of cell length (Figure S4H).

We assessed the spatial distribution of Cdr2p-mEOS3.2 detections within nodes with unitary numbers of detections (<55 detections per node) by measuring the radial distance of each detection from the center of mass of the node. The numbers of detections declined radially from the center, with 75% within 51 nm of the center (Figure 3F). All of these unitary nodes were similar in size (Figure S4 J-M).

The local densities (Figure S6) of nodes imaged around the equator by FPALM varied from cell to cell but increased modestly during interphase (Figure 3D). This is consistent with previous measurements by confocal microscopy which showed that the numbers of nodes (Deng and Moseley, 2013; Pan *et al*., 2014) increased during interphase faster than the increase in the width of the broad band of nodes (Bhatia *et al*., 2014; Pan *et al*., 2014).

### Super resolution microscopy of type 2 interphase nodes

Blt1p-mEOS3.2 appeared in discrete, punctate structures less than 100 nm in diameter near the plasma membrane throughout the cell cycle (Figure 4A, E). The appearance of these nodes was similar in side views (Figure 4, A B E and F) and face views (Figure 4G). However, clusters of detections of Blt1p-mEOS3.2 in type 2 nodes were larger near the cell poles than around the equator (Figure 4F). Temporal color coding showed that some clusters of Blt1p-mEOS3.2 moved on a scale of hundreds of nm during data acquisition giving rise to streaks or rainbow colors. Such motions are expected from the high diffusion coefficients of type 2 nodes at the tips of cells (Akamatsu *et al*., 2014).

**Figure 4.**
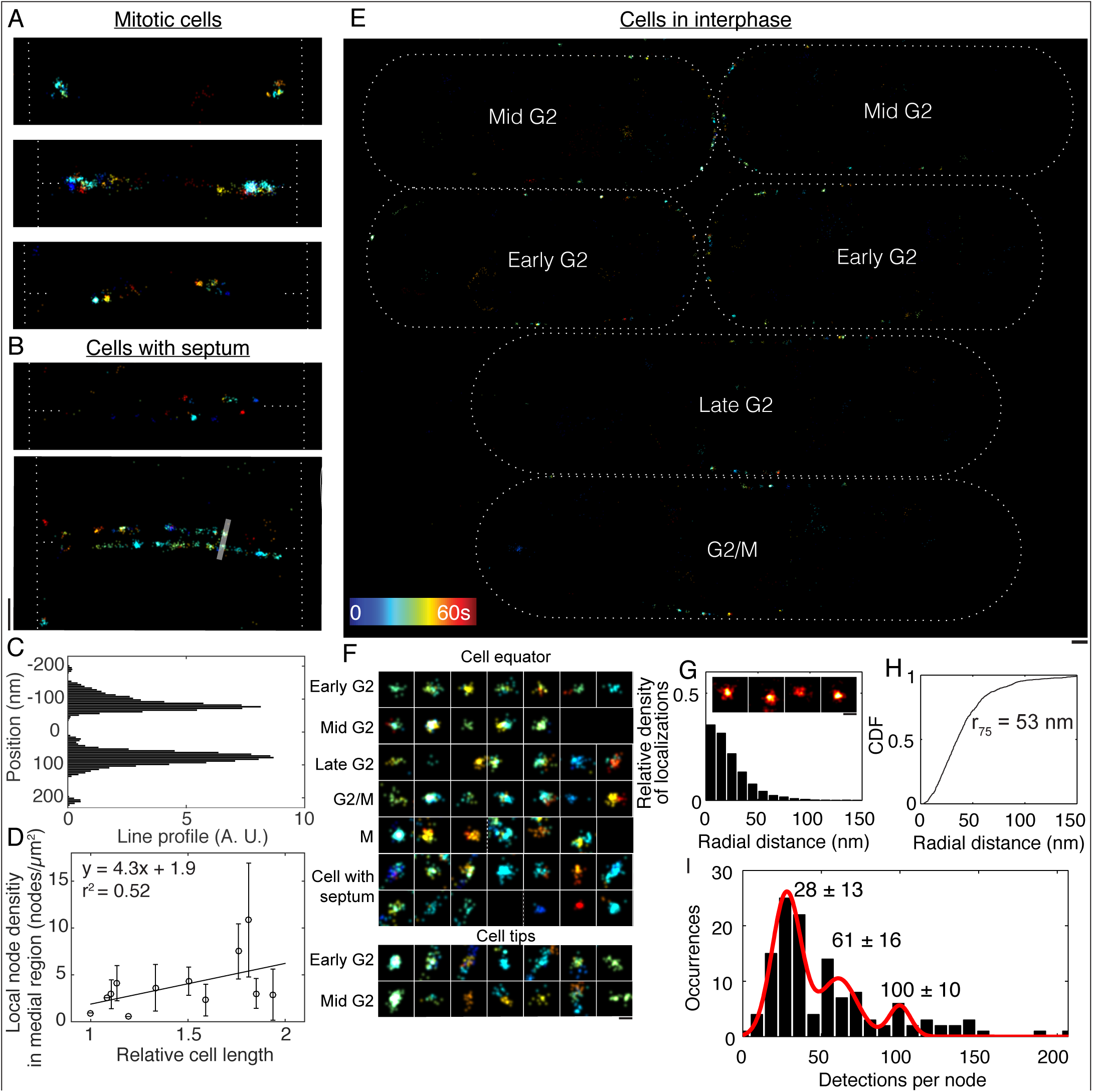
High speed FPALM of cells expressing Blt1p-mEOS3.2. A, B, E and F are displayed as Gaussian kernel time colored maps according to the times when detections occurred during acquisition. Dotted white lines mark cell perimeters and sites of division. (A) Nodes in cells with contractile rings but no septum. (B) Nodes in cells with septa. Bar is 400 nm for A and B. (C) Line scan of two nodes marked in the lower panel of (B). Intensity values (proportional to density of detections) were averaged across the 20 pixel width of the line. (D) Local surface densities of interphase nodes in a zone 1.6 μm wide centered on the equator across the cell cycle from a sample of 472 nodes in 13 cells in two fields. Line is a linear fit. (E) Image of 6 interphase cells marked with cell cycle stage. Bar is 400 nm. (F) Images of individual nodes from A, B and E, arranged by cell cycle stage and location at cell equators or cell tips. Dashed white lines separate nodes from different cells. Bar is 100 nm. (G and H) Analysis of the spatial distribution of Blt1p-mEOS3.2 in face views of nodes with fewer than 50 detections (approximately the n = 1 peak in I). (G) Histograms of the radial density distribution of detections from the center of each node. Inset: Gaussian kernel density heat maps of detections in individual nodes (face views). Bar is 100 nm. (H) Cumulative distribution plots of the radial density of detections in nodes marked by Blt1p-mEOS3.2. CDF, cumulative distribution function. (I) Histogram of the numbers of FPALM detections per node for face views of nodes. The continuous curves are fits of multiple Gaussian distributions to the data with the peak numbers of detections indicated. The 75th percentile of detection radial distances is reported in the figure. Values reported are means ± SD from the fits. n = 125 spots.

FPALM showed for the first time that Blt1p forms discrete structures in contractile rings during cytokinesis (Figure 4A). These clusters of Blt1p-mEOS3.2 in contractile rings were sometimes blurred, but were generally indistinguishable in size and shape from interphase type 2 nodes (Figure 4, A and E). These Blt1p nodes are spaced too closely in contractile rings to be resolved by confocal microscopy.

Once a dividing cell had constricted its contractile ring and formed a septum in the cleavage furrow, Blt1p-mEOS3.2 remained in two parallel disks of discrete, punctate structures spread along the plasma membrane forming the cleavage furrow (Figures 4B and S5H). These arrays of type 2 nodes were separated by ∼155 nm (Figure 4C), corresponding to the thickness of the septum (Johnson *et al*., 1973; Cortés *et al*., 2012), a distance well within the precision of the FPALM measurement (Figure S5C). These data confirm that type 2 nodes emerge from the disassembling contractile ring (Akamatsu *et al*., 2014) and associate with the newly expanded plasma membrane lining the cleavage furrow.

After the two daughter cells separated, type 2 nodes could be observed as discrete structures at the new cell poles formed by cytokinesis and also spread along the sides of the cells (Figure 4E). These nodes were blurred in the FPALM images (Figure 4F), due to the diffusion coefficients ranging from tens to ∼1800 nm^2^/s (Akamatsu *et al*., 2014).

During interphase, the type 2 node markers accumulated around the equator in discrete structures <100 nm in diameter, indistinguishable from interphase nodes elsewhere in the cell. Thus the number of type 2 nodes per unit area around the equator increased >3 fold (Figure 4D). The positions of each of these objects varied little over the course of 1 min of imaging, which leads to the conclusion that type 2 nodes around the equator were relatively stationary. This behavior is consistent with them being colocalized with and anchored by type 1 nodes at the equator during late interphase (Akamatsu *et al*., 2014).

Similar to Cdr2p-mEOS3.2 in type 1 nodes, the numbers of detections of Blt1p-mEOS3.2 per node covered a wide range but distributed into multiples of a unitary number of 28 localizations per node (Figures 4I and S4I; Table S3). Fifty-four percent of type 2 nodes marked with Blt1p were unitary, 26% were binary and the remainder had more detections. Radial density distributions of the numbers of detections fell off with distance from the center with 75% of the detections appearing within 53 nm of the centers (Figure 4, G-H).

We conclude that both type 1 and type 2 nodes are discrete structures with defined sizes and numbers of constituent proteins. While type 1 node proteins cycle between a diffuse cytoplasmic pool (during mitosis) and cortical nodes (during interphase), type 2 nodes remain intact throughout the cell cycle.

## DISCUSSION

Our quantitative measurements by confocal microscopy and FPALM support our hypothesis that both types of nodes have defined sizes and numbers of constituent proteins. These unitary type 1 and 2 nodes are relatively uniform over the cell cycle in terms of dimensions and appearance (measured by fluorescence intensity or numbers of detections) (Figure S4). Nodes appear heterogeneous by confocal microscopy, because unitary nodes cannot be resolved in local clusters of two or more nodes.

### Numbers of nodes

Three groups measured the numbers of interphase nodes by confocal microscopy. Pan et al. (2014) reported that Cdr2p-GFP nodes increased in number from 25 to 40, an acknowledged underestimate. Bhatia et al. (2015) studied the mid-focal plane during interphase where the number of nodes marked with Cdr2-GFP doubled from 10 to 20, but they did not report the total number of nodes. Deng and Moseley (2014) collected the most extensive measurements of the number of Cdr2p nodes, which increased linearly from ∼52 to 106 as cells grew longer during interphase. Our estimates are similar.

We used Deng’s counts of nodes vs. cell length (personal communication) to calculate the numbers of unitary nodes. Our FPALM observations show that the wide range of intensities of Cdr2p nodes in confocal micrographs comes from many confocal spots containing multiple unitary nodes (Fig. 2AB). About 57% of confocal “nodes” are unitary, 28% have two unitary nodes, and 15% have three or more unitary nodes. Given these ratios, the number of unitary type 1 nodes is ∼80 in early G2 and ∼160 at the end of interphase. This number of unitary type 1 nodes at the end of interphase is similar to the ∼140 unitary cytokinesis nodes that Laplante et al. (2016) measured by FPALM, given that 75-80% of type 1 nodes associate with type 2 nodes (Akamatsu *et al*., 2014) and presumably mature to cytokinesis nodes.

### Compositions of interphase nodes

We used this new estimate of node numbers and our measurements of the numbers of molecules in broad band of nodes to calculate (see methods) the approximate numbers of molecules in unitary interphase nodes at the end of G2 phase (Table 1). Like cytokinesis nodes (Laplante *et al*., 2016b) mature type 1 nodes each have about 10 molecules of Mid1p. Our estimates of 10 Mid1p molecules and 30 Cdr2p molecules per node are lower than previous measurements of Cdr2p (Pan *et al*., 2014) and Mid1p (Laporte *et al*., 2011) by confocal microscopy, because the diffraction-limited spots in those studies were not always unitary nodes. Isolated Blt1p forms tetramers (Goss *et al*., 2014), so the 20 Blt1p molecules may associate with single copies of Klp8p. Gef2p and Nod1p are also present in stoichiometric ratios (Zhu et al., 2013).

Cdr2p and Blt1p are the leading candidates to be the core proteins of type 1 and 2 interphase nodes. They are the most abundant proteins in their respective nodes and they are required for other node proteins to assemble (Almonacid *et al*., 2009; Martin and Berthelot-Grosjean, 2009; Moseley *et al*., 2009).

Both types of interphase nodes are closely associated with the plasma membrane. Multiple candidates are available to link interphase nodes to plasma membrane lipids: in type 1 nodes Cdr2p has a C-terminal KA1 domain (Moravcevic *et al*., 2010; Rincon *et al*., 2014) and Mid1p has both PH and C2 domains (Celton-Morizur *et al*., 2004; Sun *et al*., 2015) while Blt1p in type 2 nodes binds phospholipids (Guzman-Vendrell *et al*., 2013).

### Life cycle of type 1 nodes

The numbers of each interphase node protein are constant during mitosis (Carpy, 2014), but during interphase each accumulates in proportion to cell volume. In parallel, the numbers of molecules of type 1 node proteins increase around the middle of the cell due to an increase in the number of nodes, as the numbers of molecules per node (proportional to intensity or number of detections) do not change over the cell cycle (Figure S4, Bhatia *et al*., 2014; Pan *et al*., 2014).

Therefore, the biochemical state of the cell rather than gene expression controls the assembly of nodes. For example, type 1 node proteins cycle between a diffuse cytoplasmic pool (during mitosis) and cortical nodes (during interphase). The septation initiation network (SIN) signaling pathway disperses type 1 node proteins into the cytoplasm during mitosis (Morrell *et al*., 2004; Martin and Berthelot-Grosjean, 2009; Moseley *et al*., 2009; Akamatsu *et al*., 2014; Pu *et al*., 2015; Figure S3), and the decline of SIN activity allows type 1 nodes to reassemble around the equators of the daughter cells at the end of mitosis (Pu *et al*., 2015). The proteins recycle, since Cdr2p nodes remain in cells treated with the protein synthesis inhibitor cyclohexamide (Pan *et al*., 2014).

We find that type 1 nodes assemble in two phases: a burst at the end of mitosis assembles about half of these nodes, followed by steady increase in numbers during interphase to double the initial number. Type 1 nodes appear quickly at the end of the eclipse phase (Figure 3), either assembling on the plasma membrane or forming in the cytoplasm before binding to the plasma membrane. Pan and Chang (2014) proposed that type 1 nodes grow by accretion of smaller Cdr2p assemblies moving along the inner surface of the plasma membrane. However, such fast moving, small units cannot be detected by live-cell FPALM, since they generate too few single-molecule detections.

Mid1p accumulates in type 1 nodes that then mature by merging with type 2 nodes prior to accumulating cytokinesis proteins. Thus the mechanisms concentrating stationary type 1 nodes around the equator determine the distribution of type 2 nodes and site of the cleavage furrow. Mid1p comes along as a passenger on type 1 nodes rather than determining their location. In addition, Mid1p has an uncharacterized influence on the location of cytokinesis nodes, since they are not confined to the equator in *mid1Δ* cells (Daga and Chang, 2005; Wu *et al*., 2006; Almonacid *et al*., 2009; Laporte *et al*., 2011; Lee and Wu, 2012; Saha and Pollard, 2012).

### Life cycle of type 2 nodes

Type 2 nodes remain intact throughout the cell cycle including time in the contractile ring. Confocal microscopy established that Blt1p is associated with cytokinesis structures across the entire cell cycle, including interphase nodes, cytokinesis nodes and the contractile ring (Moseley *et al*., 2009; Akamatsu *et al*., 2014; Goss *et al*., 2014). However, the discrete node structure is lost to view during mitosis when the fluorescence in confocal images is spread uniformly throughout the contractile ring. This raised a question about the organization of Blt1p in the contractile ring. The ten-fold improvement in resolution provided by FPALM has provided a clear answer.

Blt1p remains in compact foci the size of interphase nodes within the contractile ring. The shape and size of these Blt1p loci are similar to those of cytokinesis nodes in contractile rings (Laplante *et al*., 2016b). As the ring constricts and disassembles, Blt1p nodes remain along the cleavage furrow and can be resolved by FPALM (Figures 4 and S5). Once the nodes spread away from the division site they can again be resolved by confocal microscopy.

### Formation of cytokinesis nodes

Cytokinesis nodes form by a series of reactions. First, diffusing type 2 nodes encounter and merge with stationary type 1 nodes, which establish the location of cleavage as shown by mutations that alter the distribution of type 2 nodes (Celton-Morizur *et al*., 2006; Padte *et al*., 2006; Moseley *et al*., 2009; Lee and Wu, 2012). The unitary nature of interphase nodes suggests that cytokinesis nodes form by docking of fully formed type 1 and type 2 nodes, a hypothesis for testing by further experimentation. Less likely, type 2 node proteins may add to type 1 nodes by accretion, as observed for Myo2 adding to cytokinesis nodes (Vavylonis *et al*., 2008). The ∼35 nm resolution of our FPALM images did not allow us to test this docking hypothesis or resolve any substructure.

Second, Mid1p transfers from type 1 to type 2 nodes before the type 1 node proteins disperse during mitosis (Morrell *et al*., 2004; Martin and Berthelot-Grosjean, 2009; Moseley *et al*., 2009; Saha and Pollard, 2012; Guzman-Vendrell *et al*., 2013; Jourdain *et al*., 2013; Zhu *et al*., 2013; Akamatsu *et al*., 2014; Goss *et al*., 2014). Gef2p and binding partner Nod1p likely aid in the transfer, as Gef2p immunoprecipitates with Mid1p and the three proteins reside stoichiometrically in nodes (Ye *et al*., 2012; Guzman-Vendrell *et al*., 2013; Zhu *et al*., 2013). Then, cytokinetic nodes slowly accumulate other cytokinesis proteins resulting in ratios of one Mid1p molecule for each myosin-II Myo2 dimer, F-BAR protein Cdc15p dimer and IQGAP Rng2p dimer (Wu and Pollard, 2005; Laporte *et al*., 2011; Laplante *et al*., 2016b). Based on the total numbers of these molecules around the equator and the numbers of unitary nodes counted by FPALM, cytokinesis nodes have about ten copies of these units. Finally, forces produced by myosin on actin filaments condense cytokinesis nodes into the contractile ring (Vavylonis *et al*., 2008).

Type 2 nodes containing Blt1p are platforms for assembly of cytokinesis nodes and ultimately the contractile ring. They are the only component of the contractile ring that remains intact throughout the cell cycle, so it is remarkable that cells lacking Blt1p are nearly normal with only modest delays in the constriction of the contractile ring (Moseley *et al*., 2009; Zhu *et al*., 2013; Goss *et al*., 2014). This indicates that cells have a reliable mechanism to back up the connections that Blt1p normally makes between proteins and the plasma membrane. For example, if mutations *cdr2Δ* or *blt1Δ* disrupt the mechanisms that normally target proteins to interphase nodes, the F-BAR protein Cdc15p slowly recruits node proteins including Gef2p and Nod1p to the contractile ring around or after the time of SPB separation (Ye *et al*., 2012; Zhu *et al*., 2013). This interdependence of redundant pathways explains the strong negative genetic interaction between mutations of *cdc15* and *blt1* (Goss *et al*., 2014). Type 2 interphase node proteins may be some of many proteins that recruit and retain Mid1p, perhaps in relationship to its phosphorylation status during mitosis.

Fits of our histograms of fluorescence intensity or FPALM detections per node suggest that interphase nodes are discrete macromolecular assemblies (quantized distributions) rather than amorphous aggregates of proteins (continuous log-normal distributions) (Figure S2, H-N; Figure S5, I-J; Table S3). A unitary structure for nodes is consistent with the stoichiometric ratios of proteins in nodes (Fig. S2G; Table 1; Coffman *et al*., 2011; Laporte *et al*., 2011; Zhu *et al*., 2013), the narrow distribution of node sizes (Figures 3E and 4H; Laplante *et al*., 2016b), and defined positions of proteins within nodes (Laporte *et al*., 2011; Laplante *et al*., 2016b). This knowledge paves the way for studies of the internal organizations of interphase nodes and their functions in cytokinesis.

### Materials and methods

#### Strain construction

We created strains of *S. pombe* with genetically encoded fluorescent proteins using standard PCR-based gene targeting methods (Bähler *et al*., 1998; Laplante *et al*., 2016a) using plasmids PFA6a-mEGFP-kanMX6 PFA6a-3GFP and pFA6a-mEos3.2-kanMX6 to insert genes for fluorescent proteins mEGFP 3GFP or mEOS3.2 upstream or downstream of the open reading frame in the endogenous chromosomal locus. We constructed the other strains by genetic crosses to laboratory stock strains.

Some fluorescent fusion proteins were not fully functional in cells. *cdr1-3GFP* cells were shorter than wild type cells (Martin and Berthelot-Grosjean, 2009; Moseley *et al*., 2009; Akamatsu *et al*., 2014) and *GFP-wee1* cells were longer than wild type cells (Moseley *et al*., 2009) Figure S1B. The additional promoter in the kanMX6 cassette increased the cellular expression of N-terminally labeled mEGFP-Gef2 by ∼3-4x (Zhu *et al*., 2013). The addition of mEOS3.2 to the C-terminus of Cdr2p or Blt1p did not affect cell growth (Figure S1B).

#### Confocal microscopy

We imaged cells on gelatin pads in growth chambers containing EMM5S media and 100 μM n-propyl gallate at 25°C. We used an inverted microscope (Olympus, IX-71) with a 100X, 1.4 NA Plan Apochromat objective (Olympus), fitted with a spinning disk confocal head (CSU-X1, Yokogawa Corporation of America), electron multiplying charge-coupled device camera (iXon 897, Andor Technology), argon-ion lasers (Melles Griot), acousto-optical tunable filters (Gooch and Housego), and dichroic mirrors and filters (Semrock). Images were acquired using Andor IQ2 software.

For still images we took Z series of 21 260 nm confocal slices encompassing 5.2 μm, which covered the entire cell. For time lapse imaging we took Z series of 3 400 nm confocal slices closest to the coverslip at 1 or 2 s intervals for ∼200 s.

#### Data analysis

*Image correction*. We corrected for uneven illumination and camera offset in the confocal micrographs by imaging slides of purified mYFP and using automated image correction software (McCormick *et al*., 2013). We corrected for uneven illumination in three dimensions, which also corrects for the ∼40% change in fluorescence intensity from the top to the bottom of the 3.5 μm diameter cells due to spherical aberrations, the difference in refractive index between the coverslip and sample. After correction the fluorescence intensity per node was similar in nodes at the top and bottom of the cell.

We wrote custom image analysis software in ImageJ64 and MATLAB (R2015a) to semi-automatically make measurements and perform calculations on the confocal stacks of images using user-defined regions of interest. ImageJ software was written using JEdit (1.0) and some functions were adapted from published software (McCormick *et al*., 2013; Akamatsu *et al*., 2014).

We corrected the final fluorescence intensity values based on the fluorescent protein used: mEGFP is 1.1x brighter than GFP (Coffman *et al*., 2011) and 3xGFP is 3 times brighter than GFP (Wu and Pollard, 2005).

For the high copy-number calibration strain *fim1-mEGFP* we reduced the electron multiplication (EM) gain from 300 to 100 to maintain the camera intensity values within the linear range, and scaled the resultant values accordingly. The EM gain was linear up to a value of 300.

*Analysis of asynchronous cells*. We used cell length as a proxy for cell cycle time (Baumgärtner and Tolic-Nørrelykke, 2009; Akamatsu *et al*., 2014). We subdivided the 4-5 h of interphase into five stages based cell morphology or length: early G2 cells were <8.5 μm long, mid-G2 cells were 8.5-10.5 μm long, late G2 cells were 10.6-12.5 μm long, and G2/M cells were >12.5 μm long. Cdr1p-3GFP cells are shorter than wild-type cells (Martin and Berthelot-Grosjean, 2009; Moseley *et al*., 2009) so we defined these stages as for wild type minus 1.5 μm (Akamatsu *et al*., 2014). Wee1p-GFP and GFP-Wee1p cells are longer than wild type cells, so we define these stages as for wild type plus 5.5 μm. For Figures 1, 3, and 4 we report the relative length of cells, defined as the difference ratio between the current cell length and the shortest cell in the population, divided by the difference between the longest and shortest cells in the population. The values range from 1 at cell birth to 2 during mitosis. The Y-intercept of these plots at x = 1 provides a measure of the mean number of molecules in the region of interest at cell birth. We scaled the length values for Cdr1p-3GFP cells independently because their mean length differs from wild type cells.

*Calculating cellular concentrations*. We counted molecules using a calibration curve that related fluorescence intensity to number of molecules based on quantitative immunoblots (Wu and Pollard, 2005; McCormick *et al*., 2013). This calibration method agrees with orthogonal methods of measuring molecules within ∼30% (Coffman *et al*., 2011; Lawrimore *et al*., 2011; McCormick *et al*., 2013). We circled cells manually and measured the fluorescence summed over all 21 260 nm slices. From the cellular regions we extracted the cell length (Feret’s diameter, defined as the longest distance between two points in the region), area, and sum fluorescence intensity for analysis. We estimated the cytoplasmic volume for each cell based on its area, the average area of the cells measured, and the assumption that the average fission yeast cell cytoplasm has a volume of 27 μm^3^ (Wu and Pollard, 2005). These concentration measurements are the total number of molecules per cell divided by the scaled estimate of cellular volume.

*Calculating number of molecules per region*. We measured the fluorescence in a region the width of the cell and containing >95% of the fluorescence of the object of interest: 1 μm wide for the contractile ring and 4 μm wide for the equatorial band of nodes. To account for cytoplasmic and cellular background we measured fluorescence from a region containing the original measurement region that was 2x greater in area, or the length of the cell if 2x the original region exceeded the cell length (Wu *et al*., 2008). For cells in early interphase with nodes near the cell tips, this background measurement did not cover the nodes concentrated at the new cell tip, but sometimes included other nodes scattered outside the broad band of nodes. We could not measure the number of molecules of Wee1p per broad band due to the low signal of GFP-Wee1p in nodes relative to background and its localization to the spindle pole body. The fraction of proteins in broad band (Figure 1, E and F) is defined as the number of molecules per broad band divided by the number of molecules per cell, on a per-cell basis. We used a two-sample Kolmogorov-Smirnov test to evaluate the significance of the increase in the fraction of molecules per broad band between early and mid G2 (Figure 1F).

*Calculating fluorescence intensity per node*. We measured the fluorescence intensity per node as the sum of fluorescence intensity in 5 consecutive 260 nm confocal slices closest to the coverslip. We made measurements of nodes containing fluorescence in at least 3 consecutive slices, and that did not have fluorescence in the adjacent slices outside the 5-slice stack. We measured background fluorescence intensity from cytoplasmic regions in the cell (Sirotkin *et al*., 2010; Coffman and Wu, 2012). For Figure S2, C-E we measured the total fluorescence within continuous regions corresponding to a node or a clump of nodes, as depicted in Figure S2A. This included regions that contained additional fluorescence outside the 5-slice stack used for quantification. For Figure S2, F and G we compared the fluorescence intensity of nodes marked by Cdr2p-mCherry and Cdr1p-3GFP in the same cell. We generated regions of interest surrounding diffraction-limited spots that contained the fluorescence of nodes in each channel. We report the intensity in each channel subtracted by the mean cytoplasmic background fluorescence in the field. To generate the background fluorescence measurements in Figure 2, we measured the fluorescence intensity for each background region of interest, subtracted by all the other background regions in that cell. We plot the mean + standard deviation of this background fluorescence value.

*Fitting Gaussian distributions to histograms*. We used the MATLAB curve fitting tool to fit each histogram in Figures 2, 3G, and 4I using a nonlinear least squares method. We used equations for the sum of three or four Gaussian distributions. We set the amplitudes, standard deviations, and means for each of the Gaussian distributions as free parameters, with constraints that the parameters be positive. We set parameter starting values for the fit as integers within the order of magnitude observed for each distribution.

*Comparison of fit quality*. We calculated the F statistic to compare the quality of the fits between multiple Gaussian distributions and a log-normal distribution for each histogram in Figures 2, 3G and 4I (Figures S2 and S4; Table S3). The F statistic is the ratio of the sum of squared errors between the two fits, normalized by the number of degrees of freedom (where a higher F value corresponds to a better fit for the first model).

*Analysis of time lapse images*. To analyze time-lapse fluorescence images of nodes we used a simplified version of software that tracks actin patches (Berro and Pollard, 2014). Regions were selected manually and then automatically re-centered at each time point with a custom algorithm using a Gaussian kernel (Berro and Pollard, 2014) and then plotted with custom MATLAB software.

For nodes that changed in fluorescence intensity over the course of imaging, we checked that the maximum fluorescence stayed in the middle slice for the duration of the imaging. We measured the background fluorescence over time from 10-20 peripheral regions containing no discrete fluorescence. We averaged the fluorescence measurements over time and between regions within a cell to generate a background measurement for each cell. The value of the background varied by 6-8% in regions within the cell (coefficient of variation 0.05-0.08) and varied by 5% between cells in an asynchronous population. We previously used these time lapse images to report the diffusion coefficients of nodes using their positions over time (Akamatsu *et al*., 2014) but we did not report their fluorescence intensities over time.

*Analysis of node number and estimate of molecules per node*. We used raw unbinned data from Deng and Moseley (2013) to calculate the number of unitary type 1 nodes. Their measurements of type 1 node number by confocal microscopy are the most reliable and agree with our own estimates by FPALM, using average surface density measurements and assumptions about the surface area of the broad band of nodes. We used their linear regression to estimate that the measured number of nodes increased from 52 (7.5 μm cells) to 106 (14.8 μm cells). We used the ratios of unitary versus multiple nodes from Fig. 2A to convert this number to the number of unitary nodes. We divided our measurements of the numbers of molecules per broad band by these measurements of node number to make estimates of the numbers of molecules per node.

### Fluorescence photoactivation localization microscopy (FPALM)

We imaged cells on gelatin pads in growth chambers containing EMM5S media and 100 μM n-propyl gallate at 25°C. We imaged cells with a custom-built single-molecule switching microscope (Laplante *et al*., 2016a) with the following modifications: fluorescence emission from mEos3.2 was collected by the objective and separated from the excitation light by a dichroic mirror (FF01-408/504/581/667/762, Semrock) and a bandpass filter (ET605/70, Chroma) before being focused onto the sensor chip of the sCMOS camera.

*FPALM acquisition*. We illuminated the cells with 3 - 8 kW/cm^2^ from the 561 nm laser for 10 s prior to acquisition to bleach autofluorescence and mEOS3.2 that had been photoconverted prior to the beginning of the experiment. In most experiments we increased the intensity of the 405 nm photoconversion laser from 0 to 38 W/cm^2^ in a series of even steps every 5 s or 15 s. To minimize premature photoactivation and bleaching we found the correct focal plane by bright field microscopy. We focused on the middle of a field of cells in bright field to image cells in a longitudinal section ∼400 nm thick. To image the surface of cells closest to the coverslip we imaged regions 1.6 μm closer to the coverslip than the medial section.

### Software used to construct super resolution images

We performed data analysis and super-resolution image reconstruction using sCMOS-specific single molecule localization algorithms (Huang *et al*., 2013). Briefly, we first corrected the raw data frames for the camera offset and gain. After single molecule identification from the calibrated images, we cropped images of the molecules (7 × 7 pixels) and used maximum-likelihood estimation to obtain the position, photon number and background information. We used a rejection algorithm based on the log-likelihood ratio metric to remove non-convergent fits or overlapping fluorescence signals from multiple emitters. Localization uncertainty estimates were determined using Cramér-Rao Lower bound (CRLB) with the sCMOS likelihood function. To reconstruct super-resolution images, we binned the localization estimates into two-dimensional (2D) histogram images with 5.15 nm × 5.15 nm pixel size. To aid visualization, each resulting image was convolved with a 2D Gaussian kernel with σ= 1.5 pixels.

### Analysis of reconstructed FPALM nodes

*Calculating local density of nodes*. We generated Voronoi diagrams (also called Voronoi tessellations) to calculate the local density of nodes from FPALM images of cells. We used ImageJ to manually select all discrete spots in the image with regions of interest 231 nm in diameter and record the spots’ X and Y positions within each cell. We wrote a MATLAB program to record the position of each node, length of the cell and its long axis angle relative to the X axis. The program retained nodes with ≥14 detections for further analysis. The program calls the function “voronoin.m” to generate a Voronoi diagram for all of the nodes in each cell. The Voronoi diagram partitions the cell surface into polygons each containing one node. For a given node and its polygon, each point on the polygon is closer to that node than to any other node. The area of each polygon is the inverse of the local density of that node. To account for overlapping nodes, we divided the area of the polygon by integer values corresponding to the number of unitary nodes, determined from the numbers of detections (see Figures 3G and 4I). To account for cell edges, the boundary was periodic and rotated according to the cell axis angle.

For the plots in Figures 3D and 4F, we calculated the mean node density for the middle 1.8 μm of the cell’s length for consistency with confocal measurements. The slopes and r squared values in these Figures were similar if the middle region was 4 μm long. *Calculating numbers of detections per node and radial distances*. We manually selected all discrete regions in 50 s reconstructions of FPALM images and saved the regions for analysis using the algorithms described in Laplante (2016a). Briefly, for each region we made 2D histograms of the detections within the region with a pixel size of 2 nm, and identified the radial symmetry center (Parthasarathy, 2012) of each region. We fit radially symmetric 2D Gaussian distributions to each FPALM node using a maximum-likelihood method and retained nodes for further analysis based on the quality of the fit.

For nodes retained after the above filtering step we measured the numbers of detections per region and reported these values as histograms in Figures 3 and 4. We fit three Gaussian distributions as explained for the histograms of fluorescence per node to identify the number of detections corresponding to unitary, double, or triple nodes.

For nodes within the unitary number of detections (≤55 detections for Cdr2p or ≤50 detections for Blt1p) we measured the distance of each detection from the node center (obtained by the Gaussian fit) and report histograms of the radial distances, with each distance bin normalized by the value of the radius. We plot the cumulative distributions of these histograms and report the radial distance that contains at least 75% of localizations, as well as the full width half maximum, defined as 2 × the 50^th^ percentile of radial distances. These two measurements are not directly comparable because each radial distance bin occupies a different amount of area.

### Online supplemental material

## Acknowledgements

This work was supported by NIH research grants GM-026132 and GM-026338 and grant 095927/A/11/Z from the Wellcome Trust. The authors thank Jian-Qiu Wu and Hirohisa Masuda for providing strains, Chad McCormick and Julien Berro for ImageJ macros, Thomas Fai for help creating Voronoi diagrams, Charlotte Kaplan for advice in FPALM localization distribution analysis, and Fang Huang for FPALM reconstruction and analysis software. J.B. discloses financial interest in Bruker Corp. (JB) and in Hamamatsu Photonics K.K.

## Abbreviation List

A. U., arbitrary units; CDF, cumulative distribution function; FPALM, fluorescence photoactivation localization microscopy; G2, gap phase of the cell cycle; GEF, guanine nucleotide exchange factor; inf, infinite (high F value); mEGFP, monomeric enhanced GFP; n. a., not applicable; n. d., not determined; mEOS3.2, monomeric EOS version 3.2; SIN, Septation Initiation Network; SD, standard deviation; SPB, spindle pole body.

## Supplemental Figure legends

Figure S1. (A) Growth at 25, 30, or 37°C on YE5S agar plates of 5-μl aliquots of wild-type cells endogenously expressing the fluorescent fusion proteins indicated. The labels indicate the numbers of cells aliquoted. n = 88 cells (wild type) or same as molecule per cell measurements in Table 1. (B) Mean lengths (± S.D.) of asynchronous cells expressing the fluorescent fusion proteins from their endogenous loci.

Figure S2. (A) Reverse contrast confocal fluorescence micrograph of a field of cells expressing Cdr2p-mEGFP with blue circles around discrete fluorescent spots (all spots) and green circles around cytoplasmic regions selected for background. (B) Reverse contrast confocal fluorescence micrograph of a field of cells expressing GFP-Wee1p at endogenous concentrations. Scale bars 2 μm. Blue circles outline spots used for quantification in Fig. 2. (C-E) Distributions of fluorescence intensities of all the nodes and clusters of interphase nodes in cells expressing (C) Cdr2p-mEGFP (n = 487 spots in 33 cells), (D) Blt1p-mEGFP (n = 294 spots in 13 cells) and (E) Mid1p-mEGFP (n = 123 spots in 8 cells). (F-G) Correlation of node fluorescence intensities between type 1 node proteins Cdr1p-3GFP and Cdr2p-mCherry in the same cell. (F) Sum projections of 4 300 nm confocal Z slices of cells expressing Cdr1p-3GFP and Cdr2p-mCherry. Regions used for node measurements outlined in yellow. (G) Graph of Cdr1p-3GFP versus Cdr2p-mCherry fluorescence per node. Line is a linear fit with r^2^ = 0.51. (H-N) Comparison of (blue) fits of a log-normal distribution and (red) fits of multiple Gaussian distributions for the histograms in Fig. 2.

Figure S3. Fluorescence intensities of Cdr2p-mEGFP and Blt1p-mEGFP nodes over 2.5 min. (A-D) Fluorescence intensity over time of Cdr2p-mEGFP nodes in (A) early G2 cells; (B) mid G2 cells; (C) late G2 cells; (D) cell entering eclipse phase (mitosis). The fluorescence intensities of these nodes are relatively constant except for one node that lost intensity and then disappeared during the eclipse phase. (E) Fluorescence intensity over time of Blt1p-mEGFP nodes. Tracks are color coded based on cell cycle stage. The fluorescence intensities of these nodes changed little over 150 s. (A-E) Cells were imaged every 2 s (Blt1p) or 1 s (Cdr2p) for 200 s, and 3 consecutive 400 nm confocal slices were taken. Images were corrected for photobleaching using cytoplasmic background fluorescence, and fluorescence intensity was measured in each node region over time. Measured nodes were largely stationary over the 200 s imaging, except for the Cdr2p nodes during the eclipse phase (D).

Figure S4. The compositions of type 1 and type 2 nodes do not change over the cell cycle. (A-G) Fluorescence per node as a function of cell length for (A-D) type 1 nodes and (E-G) type 2 nodes. These data are reported as histograms in Fig. 2. (H, I) Number of FPALM detections per node as a function of cell length for (H) Cdr2p-mEOS3.2 and (I) Blt1p-mEOS3.2. (J-M) Relationship between node size and number of detections per node for unitary (n = 1) nodes. Node size was measured (J, K) by 75^th^ percentile of radial distance of mEOS3.2 detections from node center (see Figures 3 and 4) and (L, M) by full width half maximum (two times the 50^th^ percentile of radial distance of mEOS3.2 detections from node center). (J, L) cells expressing Cdr2p-mEOS3.2. (K, M) cells expressing Blt1p-mEOS3.2. These are data for n =1 unitary nodes (<55 detections per node).

Figure S5. FPALM image acquisition and analysis. (A) Four states of fluorescent protein mEOS3.2. 405 nm laser light (blue) converts inactive mEOS3.2 (green circle) to active mEOS3.2 (red circle); then 561 nm laser light excites the converted fluorescent protein, which emits red light. mEOS3.2 may enter a dark state (lower red circle) ∼0-9 times before irreversibly photobleaching (black circle) (Zhang *et al*., 2012). (B) Illustration of FPALM image formation from raw data. Top: raw image of a Blt1p-mEOS3.2 node made from the sum of ∼80 consecutive frames at 5 ms intervals. Lower row: Left, 2D histogram of localized detections. Intensity is proportional to number of detections within each 5.15 nm^2^ pixel. Middle, density-based image reconstruction from localized detections blurred by Gaussian distributions with standard deviation of 1.5 pixels. Right, detections color coded according to the time the fluorescence events were detected during acquisition. (C) Histogram of localization precision per detection of a representative FPALM image (see Figure 4D). Localization precision is defined as 2x the localization uncertainty (averaged in X and Y). (D) Time course of the cumulative number of FPALM detections in the field shown in Figure 4D. The power of the 405 nm conversion laser was increased in 100 steps of 0.38W/cm^2^ at 1 s intervals. (E) Time course of FPALM detections over time in the same field of cells. (F) Histogram of the numbers of FPALM detections over time in 14 232 nm^2^ regions encompassing individual or groups of nodes. Bars from each square region are color coded. (G) FPALM images of the regions used in (F). The nodes are arranged based on the average time that the fluorescence events occurred for that node. (H) FPALM image of middle focal plane of 11 cells expressing Blt1p-mEOS3.2. The power of the 405nm conversion laser increased from 0 to 38 W/cm^2^ at 15 s intervals over 60 s. Scale bar 1 μm. Upper-rightmost cell is shown in Figure 3B. White lines mark cell perimeters. (I and J) Histograms of numbers of FPALM detections per node for Blt1p or Cdr2p with fits to (blue) log-normal or (red) multiple Gaussian distributions. These histograms appear in Figures 3G and 4I.

Figure S6. Illustration of calculation of node densities by Voronoi tessellation. (A) FPALM image near the surface of a fission yeast cell expressing Blt1p-mEOS3.2. White line marks cell perimeter. Nodes are color coded according to the time each fluorescent event occurred during the 60 s acquisition (see Figure 4) Scale bar 2 μm. (B) Graph of (red) positions of nodes segmented from the cell in (A) and (blue) bounding box rotated according to the cell outline in A). All discrete spots were manually selected, and spots with ≥14 detections (red) were retained for further analysis. (C) Voronoi tessellation of the nodes in B). The Voronoi tessellation partitions the cell surface into polygons each containing one node. For a given node and its polygon, each point on the polygon is closer to that node than to any other node. The area of each polygon is the inverse of the local density of that node. To account for cell edges, the boundary was made periodic and rotated according to the cell outline in panel A. To account for overlapping nodes, we divided the area of the polygon by integer values corresponding to the number of unitary nodes, determined from the numbers of detections (see Figures 3G and 4H). (D) Graph of local node densities as a function of the distance from the cell equator for the cell in (A).

**Table S1.**
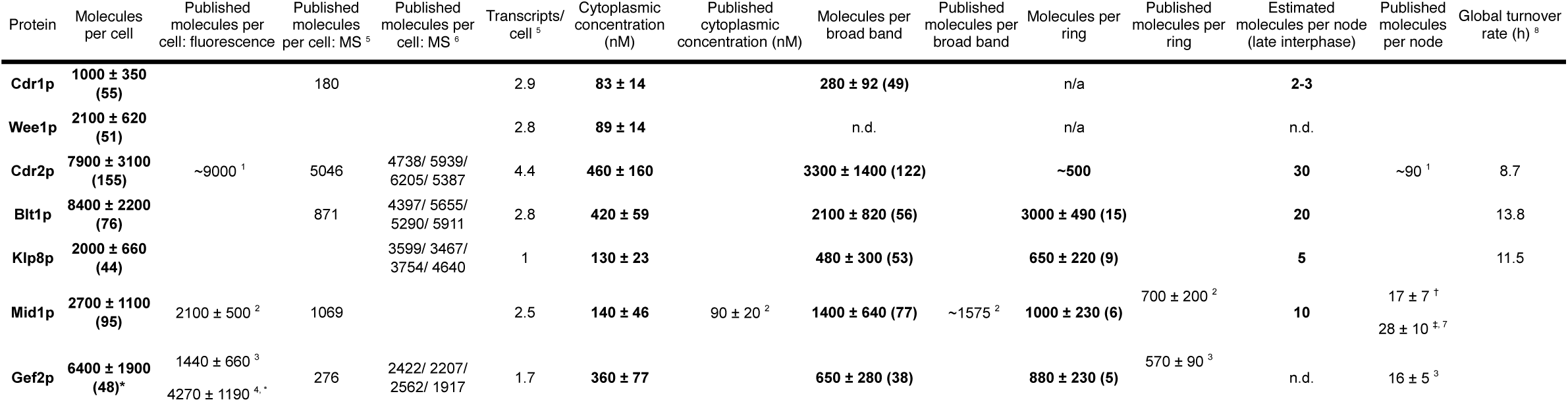
Comparison of measurements of molecule number with published values of molecule numbers and global turnover rates. Bold: measurements from this study ± s.d. with number of cells measured in parentheses. ^*^ additional promoter overexpressed protein; ^†^ interphase; ^‡^ mitosis; ^1^ Pan et al., 2014; ^2^ Wu et al., 2005; ^3^ Zhu et al., 2013; ^4^ Ye et al., 2012; ^5^ Marguerat et al., 2012 (mass spectrometry); ^6^ Carpy et al., 2014 (mass spectrometry). Measurements from *cdc25-22* cells arrested for 5 h and then released for 0, 17, 32, or 50 min. ^7^ Laporte et al., 2011 ^8^ Christiano et al., 2014.

**Table S2.**
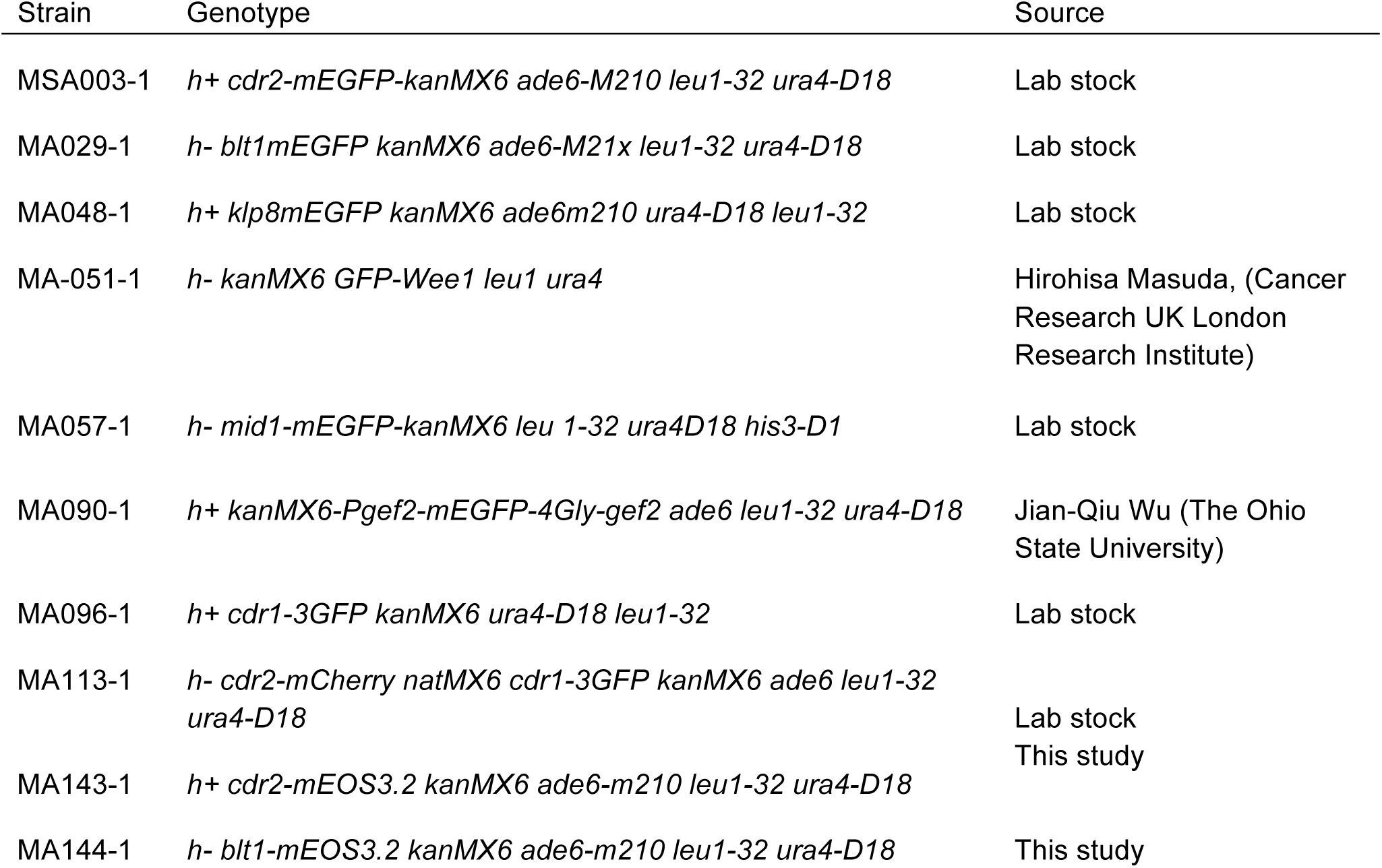
Strains used in this study.

**Table S3.** Comparison of fit quality between multiple Gaussian distributions and a log-normal distribution. Bold is better fit (higher F statistic).

| Protein | Number Gaussian distributions | F (Gaussians/ log-normal) | F (log-normal/ Gaussians) |
| --- | --- | --- | --- |
| Cdr1p-3GFP | 3 | **1.915** | 0.522 |
| GFP-Wee1p | 2 | Inf | 0.000 |
| Cdr2p-mEGFP | 4 | **21.504** | 0.047 |
| Blt1p-mEGFP | 3 | **2.885** | 0.347 |
| Klp8p-mEGFP | 2 | **1.318** | 0.759 |
| mEGFP-Gef2p | 2 | **15.640** | 0.064 |
| Mid1p-mEGFP | 2 | **2.231** | 0.448 |
| Blt1p-mEOS3.2 | 3 | **19.025** | 0.053 |
| Cdr2p-mEOS3.2 | 3 | **9.514** | 0.105 |

